# Virtually the same? Evaluating the effectiveness of remote undergraduate research experiences

**DOI:** 10.1101/2022.01.03.474815

**Authors:** Riley A. Hess, Olivia A. Erickson, Rebecca B. Cole, Jared M. Isaacs, Silvia Alvarez-Clare, Jonathan Arnold, Allison Augustus-Wallace, Joseph C. Ayoob, Alan Berkowitz, Janet Branchaw, Kevin R. Burgio, Charles H. Cannon, Ruben Michael Ceballos, C. Sarah Cohen, Hilary Coller, Jane Disney, Van A. Doze, Margaret J. Eggers, Edwin L. Ferguson, Jeffrey J. Gray, Jean T. Greenberg, Alexander Hoffmann, Danielle Jensen-Ryan, Robert M. Kao, Alex C. Keene, Johanna E. Kowalko, Steven A. Lopez, Camille Mathis, Mona Minkara, Courtney J. Murren, Mary Jo Ondrechen, Patricia Ordoñez, Anne Osano, Elizabeth Padilla-Crespo, Soubantika Palchoudhury, Hong Qin, Juan Ramírez-Lugo, Jennifer Reithel, Colin A. Shaw, Amber Smith, Rosemary J. Smith, Fern Tsien, Erin L. Dolan

**Author notes:** **Author for Correspondence:** Erin L. Dolan, University of Georgia, Biochemistry & Molecular Biology Department, B210B Davison Life Sciences Building, Athens, GA, 30602.

## Abstract

In-person undergraduate research experiences (UREs) promote students’ integration into careers in life science research. In 2020, the COVID-19 pandemic prompted institutions hosting summer URE programs to offer them remotely, raising questions about whether undergraduates who participate in remote research can experience scientific integration. To address this, we investigated indicators of scientific integration for students who participated in remote life science URE programs in summer 2020. We found that these students experienced gains in their scientific self-efficacy and scientific identity similar to results reported for in-person UREs. We also found that these students perceived high benefits and low costs of doing research at the outset of their programs, and their perceptions did not change despite the remote circumstances. Yet, their perceptions differed by program, indicating that programs differentially affected students’ perceptions of the costs of doing research. Finally, we observed that students with prior research experience made greater gains in self-efficacy and identity, as well as in their perceptions of the alignment of their values with those of the scientific community, in comparison to students with no prior research experience. This finding suggests that additional programming may be needed for undergraduates with no prior experience to benefit from remote research.

## INTRODUCTION

Undergraduate research experiences (UREs) are critical for shaping students’ decisions regarding whether to pursue graduate education and research careers in the life sciences (Gentile et al., 2017). Although UREs vary widely in duration and structure, they share some common characteristics (Gentile et al., 2017). Typically, undergraduate researchers join faculty members’ research groups to collaborate in or carry out some aspect of their research. Undergraduates are guided in their research by a more experienced researcher, such as a graduate student, postdoctoral associate, or faculty member, who is typically called their “research mentor” (Aikens et al., 2016; Joshi et al., 2019). During UREs, students are expected to engage in the practices of the discipline, including collecting and analyzing data, interpreting results, troubleshooting and problem solving, collaborating with other researchers, and communicating findings both orally and in writing (Gentile et al., 2017). Often, undergraduate researchers assume increasing ownership of their research over time, taking on greater responsibility and autonomy in their work as they gain experience and expertise.

In 2020, the COVID-19 pandemic caused massive disruptions of research, slowing or stopping research altogether at colleges and universities across the country (Korbel & Stegle, 2020; Redden, 2020). Summer URE programming was not spared from these effects. In 2019, there were 125 NSF-funded URE Sites in the biological sciences; in summer 2020, 80% of Sites were cancelled (Sally O’Conner, NSF Program Manager for BIO REU Sites, personal communication). Remarkably, about 20% of the Sites opted to proceed with their summer 2020 programs. The programs that opted to proceed were modified to operate on an entirely remote basis. Research projects had to be modified, or changed entirely, to accommodate a remote format. These modifications typically included a shift from experimental, laboratory, and field-based research and techniques to research questions or problems that could be addressed using computational and analytical approaches. Additionally, program leaders and research mentors were tasked with adapting their typical program timelines, meeting schedules, communication platforms, and curricula (e.g., seminars, workshops) to an online format.

This unprecedented and massive shift raises the question of whether undergraduates who participate in remote research programs realize the same outcomes as undergraduates who have participated in in-person research programs. This question is important to address for several reasons. First, graduate programs and employers can benefit from knowing about the experiences and outcomes of applicants whose main undergraduate research experience occurred remotely during summer 2020. Second, if remote URE programs are beneficial to students, they have the potential to dramatically expand access to research experiences, especially for students who would otherwise be excluded from in-person UREs because they have geographic constraints. Third, remote URE programs may reduce some of the cost associated with in-person programming (e.g., housing), allowing reallocation of these funds to pay additional undergraduate researchers. Finally, remote UREs may allow both students and their mentors greater flexibility in balancing work-life demands, including eliminating the hassle of relocating for a temporary summer research position. The present study aims to provide insight about whether remote UREs benefit students and thus should be considered as an option for URE programming in the future.

## THEORETICAL FRAMEWORK

For the most part, UREs have been designed to allow students to explore research as a path for further education and careers (Gentile et al., 2017; Laursen et al., 2010; Lopatto & Tobias, 2010). Multiple theories related to career development and decision-making have been used to explore and explain the outcomes students realize from participating in research. For example, Estrada, Hernandez, and colleagues carried out a series of studies framed by the Tripartite Integration Model of Social Influence (TIMSI), arguing that three social factors influence students’ integration into the scientific community (Estrada et al., 2011; Hernandez et al., 2018). Specifically, their research has shown that students’ scientific self-efficacy, scientific identity, and perceptions of the alignment between their personal values and the values of the scientific community predict whether students engage in research experiences (Estrada et al., 2011). Furthermore, students’ engagement in research increases their scientific self-efficacy, which in turn positively influences their scientific identity (Robnett et al., 2015). Thus, from an empirical perspective, research experiences can stimulate a positive feedback loop through which students develop their research skills, feel more capable of performing research, identify and share values with the research community, and choose to continue in research (Hernandez et al., 2020). Theoretically, the TIMSI illustrates how research experiences embed students in the social environment of a research group, thereby promoting their integration into the scientific community (Hernandez et al., 2020).

It is unclear whether remote research affords the same social environment for students to carry out research as an in-person experience. For example, the types of research activities that can be done at a distance are more limited, which may limit students’ development of research skills and, in turn, their scientific self-efficacy. The extent to which research mentors can provide in-the-moment guidance to help students overcome challenges is also likely to be limited because they are not working side by side. This may affect the extent to which students are successful in their research tasks, which could stymy their scientific self-efficacy development. Furthermore, students may feel less engaged in the social environment of their research group because their interactions are more time- and space-limited. This may in turn limit their feelings of being part of the research community, thereby limiting their scientific identity development. Thus, it is reasonable to question whether remote UREs would foster the same level of scientific integration as in-person UREs.

Prior research has also used Expectancy-Value Theory (EVT) (Eccles & Wigfield, 2002) as a framework for examining students’ value of UREs as a predictor of their motivation to continue in research (Ceyhan & Tillotson, 2020). Expectancy-Value Theory posits that an individual’s expectations about the degree to which they will be successful in a task (i.e., their self-efficacy) and their perceptions of the value of the task influence their motivation to engage in the task in the future (Eccles & Wigfield, 2002). From this theoretical perspective, one would expect undergraduates to decide whether to pursue graduate education or research careers based on whether they perceived they were sufficiently competent and whether doing research would provide sufficient value over costs. Value can take the forms of being personally interesting (intrinsic value), being useful (utility value), and providing prestige or respect (attainment value) (Eccles & Wigfield, 2002). Cost can be experienced in terms of effort spent, emotional or psychological tolls, or missed opportunities (Ceyhan & Tillotson, 2020).

Work from Ceyhan & Tillotson (2020) indicates that undergraduates express intrinsic and utility value as well as opportunity costs of in-person research. However, students may experience remote research differently, ascribing different values and costs to research and differing in their motivation to continue research in the future. For example, students carrying out research remotely may not be responsible for the hands-on collection of their data, which may limit their interest in the work (i.e., less intrinsic value). In contrast, students may perceive greater utility value because they learn computational skills that are useful in a variety of career paths and in high demand among employers. In addition, students may perceive less opportunity cost of doing remote research because of its inherent flexibility (e.g., no need to physically relocate, options to schedule research tasks around other personal demands).

In summary, prior research using TIMSI and EVT shows that UREs influence students’ scientific self-efficacy, scientific identity, and perceptions of the value and costs of research, which can in turn influence their intentions to pursue a graduate degree and/or a research career and their actual pursuit of these paths. Here we used these same frameworks to study of the influence of remote UREs on student outcomes.

Specifically, we sought to address the following research questions:

1. To what extent do undergraduates who engage in remote research programs experience scientific integration in terms of gains in their scientific self-efficacy, scientific identity, values alignment, and intentions to pursue graduate education and science- and research-related careers?
2. To what extent do undergraduates who engage in remote research programs shift their perceptions of the values and costs of doing research?

Due to COVID-19, it was not possible to include a comparison group of in-person undergraduate researchers. Thus, we report our results here and interpret them with respect to published results of in-person UREs, which include students in URE Sites and other URE formats (e.g., Hernandez et al., 2020; Robnett et al., 2015).

## METHODS

Here we describe the results of a single-arm, comparative study. We collected data using established survey measures of the constructs of interest, which we administered before and after students participated in a remote research program. We evaluated the measurement models, ultimately grouping values- and cost-related data into a higher order measurement model based on our results. Then we evaluated the fit of the data to a series of five multilevel random intercepts models to identify changes in our constructs of interest. The results reported here are part of a larger study of remote UREs that was reviewed and determined to be exempt by the University of Georgia Institutional Review Board (STUDY00005841, MOD00008085).

### Context and Participants

We contacted the 25 institutions that planned to host remote research programs during summer 2020 (Sally O’Connor, personal communication) to invite them to collaborate in this study. A total of 23 programs hosted by 24 research institutions in 18 states and 1 U.S. territory agreed to participate by distributing study information to their summer 2020 cohort of undergraduate researchers. The sample included 5 non-degree granting research institutes as well as 3 masters universities, 1 doctoral university, 2 high research activity universities, and 11 very high research activity universities according to the Carnegie Classification of Institutes of Higher Education. Three universities were classified as Hispanic Serving Institutions. At the time of enrollment, undergraduate researchers did not yet know that their summer programs would take place remotely. One institution did not have the capacity to host their complete program remotely, so they partnered with another institution to host a joint program. Additionally, one of the 24 institutions offered two distinct programs funded from different sources. We treated them as a single program because the participating students, their research projects, and the program activities were quite similar. In total, 307 students received the recruitment email and study information. This number includes students (n=27) who participated primarily in-person who were later excluded from the analysis. A total of 227 remote students in 22 programs (average group size=∼12) completed both the pre and postsurvey. The average program duration was ∼9 weeks; detailed duration data can be found in Table 1.

**Table 1.** Duration of URE programs. Remote URE programs in this study varied in duration, with most being about 10 weeks long. *One program had staggered end dates with most students engaging in research for 9 weeks.

| Duration in Weeks | Number of Programs |
| --- | --- |
| 5 | 1 |
| 8 | 3 |
| 9 | 4* |
| 10 | 12 |
| 11 | 2 |

The programs in this study were funded by the National Science Foundation (NSF) or the U.S. Department of Agriculture. The NSF supports UREs through two funding mechanisms: Research Experience for Undergraduate (REU) Sites, which host cohorts of students each year, or REU Supplements, which typically support one or two undergraduate researchers associated with a funded research project (National Science Foundation, n.d.). Here we focus on URE Sites, which typically offer some combination of networking with faculty and professional development to complement the mentored research experience (National Science Foundation, n.d.). In the past, URE participants have typically been junior- or senior-level undergraduate students who have committed to a STEM major, but programs are increasingly involving students at earlier points in their undergraduate career in order to attract students to a STEM career who were otherwise not interested (National Science Foundation, n.d.).

### Data Collection

We surveyed students twice using the secure survey service Qualtrics: at the beginning of their program (presurvey or Time 1) and after all program activities had been completed (postsurvey or Time 2). Students participating in programs that offered pre-program workshops were asked to complete the initial survey before engaging in these workshops. Students were sent emails with the final survey within a week of finishing their URE programs with up to two reminders. Monetary incentives were not offered. Only students who completed both surveys were included in the sample (Table 2). The survey measures are described briefly here and included in their entirety in the Supplemental Materials (Tables S1-S3).

**Table 2.** Demographics of study participants. In total, 227 students responded to both the pre- and post-survey, including 153 women, 69 men, and 4 individuals who identified as non-binary. Note that students were able to indicate multiple races or ethnicities, so race/ethnicity counts do not sum to the total sample size. With respect to parent education level, 79 students had a parent or guardian who did not attend college. There were 45 students who indicated that they had transferred to their current institution from another college or university.

| Race/Ethnicity | Previous Research Experience |  |  |  |  | Total |
| --- | --- | --- | --- | --- | --- | --- |
|  | None | 1 Term | 2 Terms | 3 Terms | >3 Terms |  |
| African American or Black | 7 | 6 | 7 | 2 | 9 | 31 |
| Central and East Asian | 6 | 5 | 8 | 7 | 4 | 30 |
| Latinx | 10 | 13 | 16 | 11 | 10 | 60 |
| Middle Eastern | - |  | 1 | - | 1 | 2 |
| Native American or Native Hawaiian | 2 | 2 | 2 | - | 1 | 7 |
| South Asian | - | 3 | 1 | - | 4 | 8 |
| White | 18 | 30 | 34 | 13 | 21 | 116 |

#### Scientific Self-Efficacy

Scientific self-efficacy is the extent to which students are confident in their ability to carry out various science research practices, such as developing a hypothesis to test. We used a 9-item Scientific Self-Efficacy measure that was a combination of 7 published items (Chemers et al., 2011; Estrada et al., 2011) and 2 items (“Use computational skills” and “Troubleshoot an investigation or experiment”) that we authored based on input from the directors of the URE programs in this study. These items were intended to more fully capture the forms of scientific self-efficacy students could develop by engaging in remote research (see Table S1 in Supplemental Materials for items). Response options ranged from 1 (“not confident”) to 6 (“extremely confident”). Responses were averaged into a single score, with higher scores indicating higher levels of scientific self-efficacy.

#### Scientific Identity

Scientific identity is the extent to which students see themselves as scientists and as members of the scientific community. We used a 7-item Scientific Identity measure using 7 published items (Chemers et al., 2011; Estrada et al., 2011) (see Table S2 in Supplemental Materials for items). An example item is “I have a strong sense of belonging to the community of scientists.” Response options ranged from 1 (“strongly disagree”) to 6 (“strongly agree”). Responses were averaged into a single score, with higher scores indicating higher levels of scientific identity.

#### Values Alignment

Science values alignment is the extent to which students see their personal values as aligning with values of the scientific community. We used a published 4-item Values Alignment measure (Estrada et al., 2011), the structure of which was based upon the Portrait Value Questionnaire (Schwartz et al., 2001) (see Table S3 in Supplemental Materials for items). Response options ranged from 1 (“not like me”) to 6 (“extremely like me”). An example item is “A person who thinks it is valuable to conduct research that builds the world’s scientific knowledge.” Responses were averaged into a single score, with higher scores indicating higher a higher degree of alignment between the student’s values and the values of the scientific community.

#### Intrinsic Value

Intrinsic value refers to how much students find research personally interesting and enjoyable. We adapted a published 6-item intrinsic value measure (Gaspard, Dicke, Flunger, Schreier, et al., 2015) (see Table S3 in Supplemental Materials for items). Response options ranged from 1 (“strongly disagree”) to 6 (“strongly agree”). An example item is “Research is fun to me.” Responses were averaged into a single score, with higher scores indicating higher levels of intrinsic value.

#### Personal Importance

Personal importance (also known as attainment value) refers to the importance that students place on doing well in research, including how relevant doing well in research is for their identity. We adapted a 3-item personal importance measure (Gaspard, Dicke, Flunger, Schreier, et al., 2015) (see Table S3 in Supplemental Materials for items). Response options ranged from 1 (“strongly disagree”) to 6 (“strongly agree”). An example item is “Research is very important to me personally.” Responses were averaged into a single score, with higher scores indicating higher levels of personal importance.

#### Utility Value

Although EVT conceptualizes utility value as a single construct, work from Gaspard and others has shown that students perceive different forms of utility from their educational experiences, such as utility for their future careers or for helping their community (Gaspard, Dicke, Flunger, Brisson, et al., 2015; Gaspard, Dicke, Flunger, Schreier, et al., 2015; Thoman et al., 2014). Thus, we chose to use a measure of utility value that included multiple dimensions: social, job, and life utility (see Table S3 in Supplemental Materials for items). Social utility refers to students’ perceptions of how useful the ability to do research would be for their communities. We adapted 3 social utility items (Gaspard, Dicke, Flunger, Schreier, et al., 2015), such as “Being well versed in research will prepare me to help my community.” Job utility refers to students’ perceptions of how useful the ability to do research would be in the context of a workplace. We adapted 3 job utility items (Gaspard, Dicke, Flunger, Schreier, et al., 2015), such as “The skills I develop in research will help me be successful in my career.” Life utility refers to students’ perceptions of how useful the ability to do research would be for their everyday lives. We adapted 3 life utility items (Gaspard, Dicke, Flunger, Schreier, et al., 2015), such as “Research comes in handy in everyday life.” For all utility items, the response options ranged from 1 (“strongly disagree”) to 6 (“strongly agree”). Responses were averaged into a single score, with higher scores indicating higher levels of utility value.

#### Cost

Cost is the extent to which students perceive research as requiring them to make sacrifices. We adapted the 3-item cost scale (Gaspard, Dicke, Flunger, Schreier, et al., 2015) (see Table S3 in Supplemental Materials for items). Response options ranged from 1 (“strongly disagree”) to 6 (“strongly agree”). An example item is “I have to give up a lot to do well in research.” Responses were averaged into a single score, with higher scores indicating higher levels of perceived cost of engaging in research.

#### Graduate and Career Intentions

Graduate and career intentions refer the extent to which students intend to pursue a graduate degree or science- or research-related career. The career-related item was used from Estrada et al. (2011) and the graduate degree related item was similarly worded, with “career” replaced with “graduate degree.” Response options ranged from 1 (“I DEFINITELY WILL NOT pursue a graduate degree in science/ a science research-related career”) and 5 (“I DEFINITELY WILL pursue a graduate degree in science/ a science research-related career”).

#### Previous Research Experience

In order to better characterize the study sample and explore possible differential effects of remote research experiences for students with different levels of research experience, we asked students how much research experience they had prior to participating in the study. Response options included: None, one semester or summer, two semesters or summers, three semesters or summers, and more than three semesters or summers.

### Data Analysis

Following the Anderson and Gerbing (1988) two-step approach, we first tested a confirmatory measurement model before fitting our structural models. Our confirmatory measurement model specifies the relationships between survey items and the latent variables they represent. Our structural models estimate the effect of participating in a remote research program on student outcomes. To attain optimum model fit for our measurement model, we followed an iterative process of model specification using confirmatory factor analysis (CFA). To test our structural model, we used a multilevel modeling approach because the data are clustered such that students are nested within programs. All analyses were conducted in R version 4.0.1 and RStudio using lme4 (linear mixed effects modeling) and lavaan (latent variable modeling) (Bates et al., 2014; Rosseel, 2012). Fixed-effect only models were estimated with maximum likelihood estimation and mixed-effect models were estimated with restricted maximum likelihood estimation, as is recommended by Theobald (2018). Conditional *R*^*2*^ values, which take into account the variance of both the fixed and random effects, were calculated using the MuMIn package for model averaging (Bartoń, 2020). Random and fixed effects for each model, as well as AIC and *R*^2^ values, are reported.

#### Assessment of Measurement Models

We used several fit indices to assess how adequately our CFA models reproduced their variance-covariance matrices. First, we report a chi squared test (*χ*^2^) for each model. Chi square is highly sensitive to misfit because it has strong assumptions, including that there is no kurtosis in the data, which is a measure of the “tailedness” of the probability distribution of a real-valued random variable (Kline, 2015). However, a significant chi square indicates misfit to some degree (Credé & Harms, 2019), and so it is best practice to report it. We also include the root mean square error of residuals (RMSEA), which approximates how well the model estimates the population covariance matrix while favoring more parsimonious models. Higher values of RMSEA indicate poorer fit. Hu and Bentler (1999) recommend an RMSEA cutoff value of 0.06. In addition, we chose to include the standardized root mean square residual (SRMSR/SRMR) because it is sensitive to mis-specified covariance structures. This means that a high SRMR value (greater than 0.08) in the absence of other indications of misfit may indicate that the factor structure is mis-specified. Finally, we consider the Tucker-Lewis index (TLI) and comparative fit index (CFI), which are incremental fit measures, meaning that they compare model fit to the worst possible model. Higher values indicate better fit. Because TLI and CFI are sensitive to mis-specified factor loadings, they are useful for evaluating the appropriateness of survey items as representative of their latent variable (Hu & Bentler, 1999). A value of 0.90 or above is recommended (Hu & Bentler, 1999).

In addition to fit indices, we evaluated the appropriateness of our measurement models based on factor loadings and coefficient alpha values (see Tables S1-S3 in Supplemental Materials for factor loadings). Factor loadings indicate the extent to which each survey item reflects its respective latent variable. A minimum factor loading of 0.40 is recommended (Bandalos, 2018). Coefficient alpha is a measure of internal consistency, or the degree or item correlation within the factor. Coefficient alpha values were similar across timepoints; we report values that include both timepoints for each measure. Ultimately, we balanced evidence from fit indices, factor loadings, and alpha values to determine our final measurement models.

##### Scientific Self-Efficacy

The scientific self-efficacy scale demonstrated high internal reliability (α=0.92). Fit of the model was acceptable, *χ*^2^ (27)=140.839 (*p<*0.001), RMSEA=0.137, SRMR=0.050, CFI=0.912, TLI=0.883, although RMSEA is substantially higher than the recommended value of 0.05 and TLI is slightly lower than the recommended value of 0.90. Given the high alpha value, the high factor loadings (0.45-0.87), and the use of this scale in the study of other UREs, we opted to proceed as is with the measure as a single factor. Item 2 (“Use computational skills [software, algorithms, and/or quantitative technologies]”) produced a factor loading much lower than the second lowest factor loading (0.45 vs. 0.66). This result suggests that students responded differently to this item. However, removing this item did not result in improved model fit, *χ*^2^ (28)=110.981 (*p<*0.001), RMSEA=0.142, SRMR=0.043, CFI=0.926, TLI=0.896. Moreover, we felt that this item captured information relevant to students’ remote research experiences. Thus, we moved forward with the complete scientific self-efficacy measure as it was administered to students.

##### Scientific Identity

The scientific identity scale also demonstrated high internal reliability (α**=**0.90). However, RMSEA, CFI, and TLI indicated poor model fit, *χ*^2^ (14)=176.429 (*p<*0.001), RMSEA=0.228, SRMR=0.096, CFI=0.792, TLI=0.688, with no clear cause of the model misfit. We attempted to remove items and test a two-factor structures with no improvement in model fit. Thus, the factor structure of scientific identity is still uncertain and may be sample dependent. Given the high alpha value, the high factor loadings (0.52-0.90), and the use of this scale in the study of other UREs, we opted to proceed with the measure as a single factor.

##### Values and Cost

We began by testing the factor structure of values with seven factors: values alignment, intrinsic value, personal importance, cost, social utility, job utility, and life utility. Overall, factor loadings were higher than the recommended minimum value of 0.40 (Bandalos, 2018), ranging from 0.473 to 0.949. Despite high factor loadings, model fit statistics indicated poor fit (*χ*^2^ (254)=747.528 (*p<0*.001), RMSEA=0.094, SRMR=0.090, TLI=0.816, CFI=0.844). Most factor correlations between the seven factors were moderate to low; however, the factor correlation between intrinsic value and personal importance was high (*r=*0.848, *p<*0.001). Therefore, we evaluated our values factor for sources of misfit. Based on item content and factor loadings in our seven-factor model, intrinsic value appeared to be two separate variables. The content of the first three items refers to enjoyment of research (e.g., “Research is fun for me”) whereas the last three items are more value-oriented (e.g., “Performing well in research is important to me”). In addition, factor loadings were stronger for the first three items (0.91, 0.95, 0.87) than for the later three items (0.60, 0.57, 0.47). The differences in the strength between the first and second half of the items suggests that the intrinsic value factor may be better represented as two factors. Indeed, when we split this factor in two, factor loadings for the second half of items (intrinsic 2) increased substantially (0.79, 0.89, 0.77), as did model fit (*χ*^2^ (247)=477.332 (*p<*0.001), RMSEA=0.065, SRMR=0.055, CFI=0.927, TLI=0.912).

##### Higher-Order Confirmatory Factor Analysis

To address concerns about measurement model fit and to simplify the interpretation of our structural model analyses, we conducted a higher-order CFA. Statistically, a “higher-order factor” models the covariance between two or more “lower-order factor(s),” which are seen as manifestations of the higher-order factor. Higher-order factors are useful because they tend to have higher predictive validity compared to narrower factors (Credé & Harms, 2015). They also help address high inter-factor correlations (Table S5). High factor correlations (*r* > 0.70) are problematic because they indicate too much overlap between constructs for them to be meaningfully different from one another. Collapsing factors into one higher-level factor addresses this concern.

Because values alignment did not correlate highly (*r* > 0.70) with any other value-related factors, we opted to represent values alignment under its own higher-order factor, HO1. Personal importance strongly correlated with intrinsic 1 and intrinsic 2, thus we chose to represent personal importance, intrinsic 1, and intrinsic 2 with a higher order factor, HO2. Because cost did not correlate strongly with other values-related factors, we represented it with the higher-order factor HO3. Finally, we group together the three forms of utility (social, job, life), based on their higher correlations and conceptual similarity, under HO4. Although the fit of this four-factor, higher-order model was good according to fit indices (*χ*^2^ (263)=525.357, *p*<0.0001, RMSEA=0.067, SRMR=0.068, CFI=0.917, TLI=0.906), there were two Heywood cases (i.e., impossible factor loadings). The standardized loading for life utility onto HO4 was 1.010 and the standardized loading for personal importance on HO2 was 1.001. Furthermore, life utility demonstrated a negative variance (−0.019), as did personal importance (−0.002) indicating misfit. HO2 was highly correlated with HO4 (*r*=0.722), so we decided to collapse the HO2 and HO4 factors to eliminate this source of misfit.

Ultimately, we fit a values and cost model that contained three higher-order factors: “Alignment” or HO1 represents students’ perceptions of values alignment, “Perceived Benefits” or HO2 represents students’ perceptions of the intrinsic value, personal importance, and utility of engaging in research, and “Perceived Costs” or HO3 represents students’ perceptions of the costs of engaging in research. For readability, we refer to these factors as alignment, perceived benefits, and perceived costs. Fit of this model was acceptable (*χ*^2^ (266)=577.278, *p*<0.0001, RMSEA=0.073, SRMR=0.080, CFI=0.902, TLI=0.889), and it eliminated the Heywood cases and negative factor variance. Thus, we decided to move forward with this three-factor model (Figure 1; see Table S4 for higher-order factor loadings and Table S5 for higher-order factor correlations).

**Figure 1.**
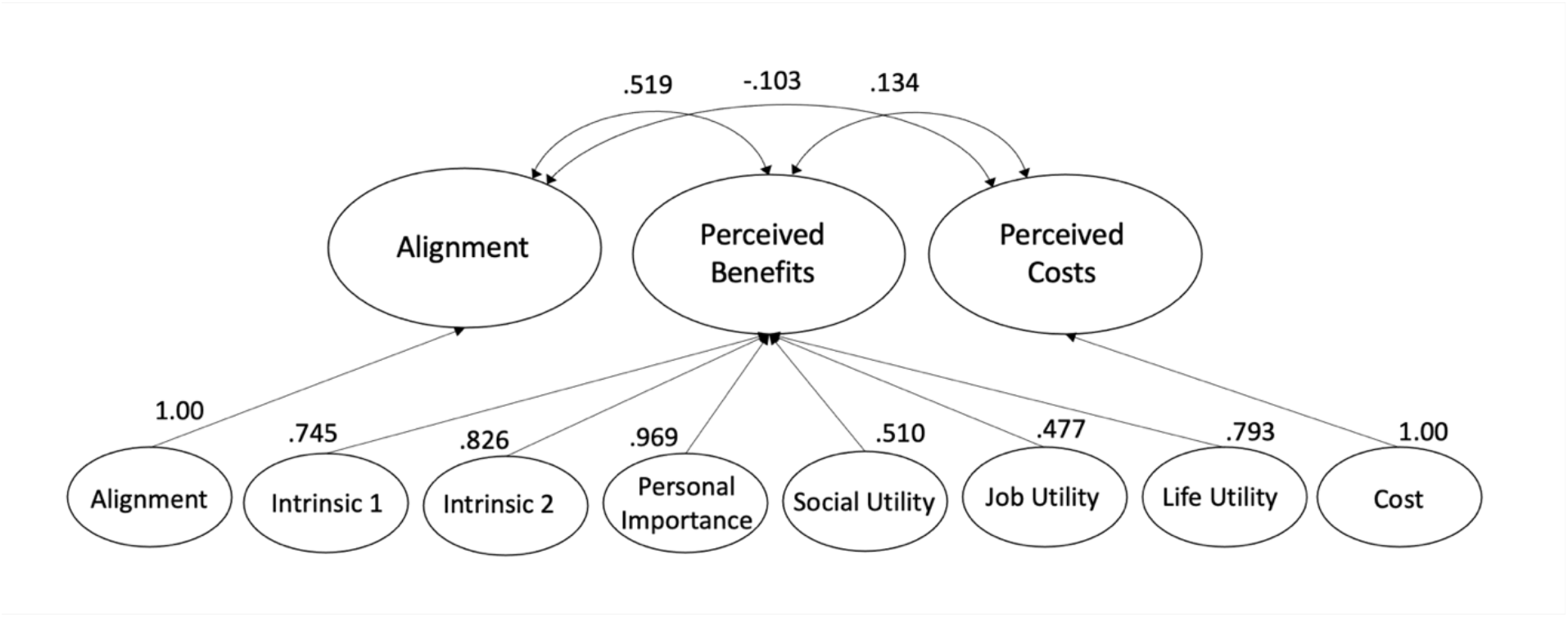
Factor loadings and factor correlations for higher-order confirmatory factor analysis. Factor loadings and correlations are reported for Time 1 (pre-URE). Loadings for the higher-order factors with only one lower-order construct (i.e., Alignment, Cost) will always be 1.00 and are not meaningful. See Supplemental Materials for Time 2 factor loadings.

#### Assessment of Structural Models

Given the exploratory nature of research on remote URE programming, we tested three models for each student outcome variable. This approach allowed us to estimate the effects of completing a remote URE to answer our research questions, and to explore whether the program in which students completed their remote URE and their level of prior research experience influenced their outcomes.

##### Model 1

This model allowed us to estimate the effects of completing a remote URE and to explore program-level effects. Specifically, there were multiple students in each program, which means that students’ experiences within programs are not independent of one another (i.e., data are clustered). Therefore, Model 1 includes program as a random effect such that each grouping factor has its own random intercept, meaning that each program’s level of our five latent variables starts at a different point on the y-axis. It also includes a fixed effect of the URE. Thus, Model 1 can be stated as:

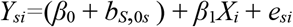

In this model, *X*_*i*_ is our predictor variable, time, which takes on a value of 0 or 1 depending on whether _*i*_ is at time 1 (pre-program) or time 2 (post-program). *e*_*si*_ represents error. *β*_0_ is the fixed effect of the slope, *β*_1_ is the fixed effect of the intercept, and *b*_S_,_0*s*_ are the random intercepts. Here we report the syntax used to run our multilevel regression models. “Student outcome variable” represents each dependent variable, “Time” represents the measurement timepoint, and “Program” represents the program where the student participated in their URE. Program is treated as a categorical variable and the student outcome variable and time are treated as continuous variables. The model syntax is as follows:

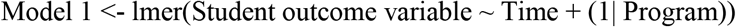

##### Model 2

Students began their UREs with different levels of research experience, which could account for variance in our dependent variables. Thus, we also included prior research experience as a fixed effect in our models. Prior research experience is treated as a categorical variable. Thus, Model 2 can be stated as:

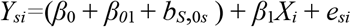

Note that this model is the same as Model 1, but with the addition of a fixed intercept for research experience, *β*_*0*1._ The model syntax may be written as:

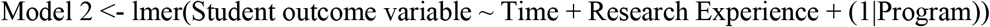

##### Model 3 Equation

Model 2 estimates the amount of variance accounted for by prior research experience. However, it does not estimate the relative importance of different levels of research experience. In other words, do more experienced researchers or less experienced researchers have more to gain from the URE? To answer this question, we estimated Model 3, which includes research experience as a random intercept. Thus, Model 3 can be stated as:

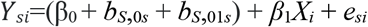

Model 3 is the same as Model 1, but with the addition of an additional random intercept of research experience, *b*_*S*_,_01*s*_. The model syntax is as follows:

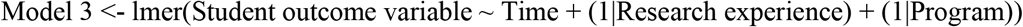

##### Comparing Models with Akaike’s Information Criteria (AIC)

In order to identify the most explanatory and parsimonious models, we chose to compare fit between models using Akaike’s information criteria (AIC). AIC is a fit index that weights how well the model fits the data, while adding a penalty for the number of parameters in the model. This penalty favors more parsimonious models, thereby balancing the likelihood function given the observations and number of parameters. Smaller AIC values indicate better-fitting models (Theobald, 2018). A difference of 2 or greater is necessary for establishing significantly different AIC values (Burnham & Anderson, 2002).

We tested a total of three models for each student outcome and compared AIC values among them. For each dependent variable, we began by testing a mixed effects model with a fixed effect of the URE and a random intercept for the program (Model 1). Next, we added in a fixed effect of students’ prior research experience (Model 2). Finally, we tested this same model with a random effect of prior research experience instead of a fixed effect (Model 3). Model 3 had the lowest AIC values and highest *R*^2^ values, and therefore is the primary model which we interpret in the following section. We also discuss the fixed effects of prior research experience from Model 2 because the fixed effects inform the strength and direction of the effect of UREs on student outcomes. Because we ran all three models seven times – once for each dependent variable – we implemented a study-wide Bonferroni correction to interpret *p*<0.007 as significant.

## RESULTS

Here we report the significant results of our Model 2 and 3 analyses (see Supplemental Materials for Model 1 and non-significant results). We report intercepts (β_0_) as a “baseline” of where students are with respect to each construct at the start of their remote URE, slopes (β_1_) to identify any changes pre- to post-URE and characterize the size of any effects, and percentages of variance in student outcomes explained by their program and their prior experience.

### Scientific Self-Efficacy

We found that students began their UREs with a moderate level of scientific self-efficacy (Model 3: β_0_=3.62, *SE=*0.07, *p*<0.001), and their self-efficacy increased significantly from pre- to post-URE (Model 3: β_1_=0.64, *SE=*0.08, *p*<0.001) (Table 3). We observed that students’ program accounted for only 3% of variance in scientific self-efficacy, which indicates that differences between programs had little if any effect on students’ self-efficacy development. We found that students’ prior research experience accounted for 9% of variance in their self-efficacy growth. Students who had three semesters or summers of prior research experience (Model 2: β=0.65, *SE*=0.15, *p*<0.0001) or more than three semesters or summers of prior research (Model 2: β=0.71, *SE*=0.13, *p<*0.0001) experienced significant gains in scientific self-efficacy. Thus, we can infer that there was a positive effect of the remote URE on students’ scientific self-efficacy, and the effect was stronger for more experienced students.

**Table 3.**
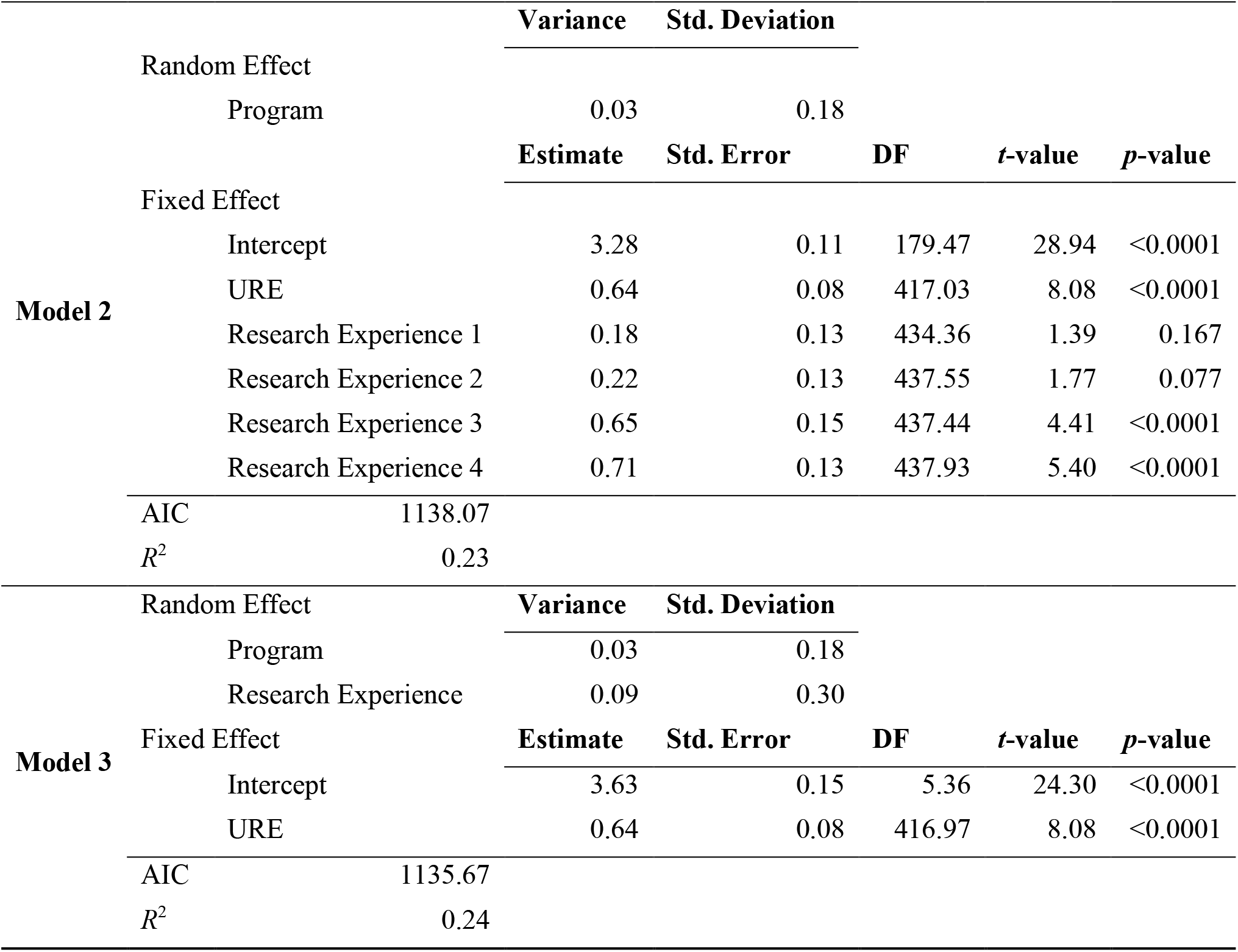
Remote UREs and prior research experience, but not program, relate to student gains in scientific self-efficacy. Students reported significantly higher levels of scientific self-efficacy from pre- to post-URE. Program had a very small effect on students’ scientific self-efficacy gains. Students with at least three semesters of prior research experience made larger gains in scientific self-efficacy compared to students with less prior experience.

In analyzing the self-efficacy data, we observed that the mean score for item 2 (“Use computational skills [software, algorithms, and/or quantitative technologies]”) is lower than for the other items in the scale: *M*=3.08 pre-URE (vs. *M=*3.42-4.10 for other items) and *M*=4.00 post-URE (vs. *M*=3.85-4.74 for other items). This suggests that, even though students are experiencing self-efficacy growth, students perceived themselves to be less capable in their computational skills.

### Scientific Identity

We found that students began their UREs at a high level of scientific identity (Model 3: β_0_*=*4.72, *SE=*0.13, *p<*0.001), which increased significantly from pre to post URE (Model 3: β_1_*=*0.24, *SE=*0.08, *p*=0.005) (Table 4). Again, we observed that the students’ program accounted for a small amount (5%) of the variance in their scientific identity growth. We also found that students’ prior research experience also accounted a small amount of the variance in scientific identity (5%), with students with more than three semesters or summers of research (Model 2: β=0.44, *p*=0.002) experiencing significantly greater gains in their sense of scientific identity.

**Table 4.**
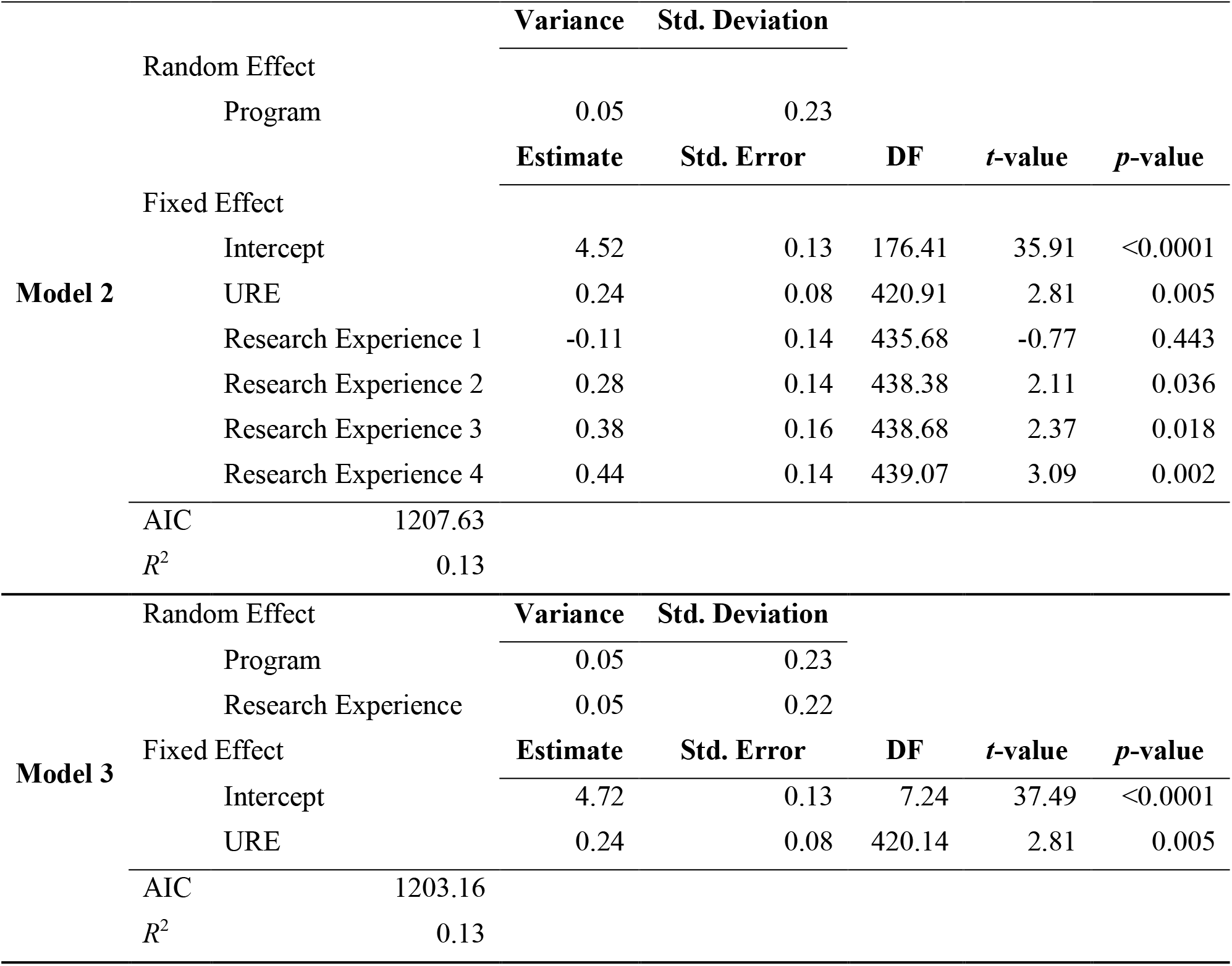
Remote UREs, program, and prior research experience relate to student gains in scientific identity. Students reported significantly higher levels of scientific identity from pre- to post-URE. Program and prior research experience had a very small effect on students’ scientific identity gains; students with at least three semesters of prior research experience made larger gains in scientific identity compared to students with less prior experience.

|  |  | Variance | Std. Deviation |  |  |  |
| --- | --- | --- | --- | --- | --- | --- |
| **Model 2** | Random Effect |  |  |  |  |  |
|  | Program | 0.05 | 0.23 |  |  |  |
|  |  | Estimate | Std. Error | DF | t-value | p-value |
|  | Fixed Effect |  |  |  |  |  |
|  | Intercept | 4.52 | 0.13 | 176.41 | 35.91 | <0.0001 |
|  | URE | 0.24 | 0.08 | 420.91 | 2.81 | 0.005 |
|  | Research Experience 1 | -0.11 | 0.14 | 435.68 | -0.77 | 0.443 |
|  | Research Experience 2 | 0.28 | 0.14 | 438.38 | 2.11 | 0.036 |
|  | Research Experience 3 | 0.38 | 0.16 | 438.68 | 2.37 | 0.018 |
|  | Research Experience 4 | 0.44 | 0.14 | 439.07 | 3.09 | 0.002 |
| AIC |  | 1207.63 |  |  |  |  |
| $R^2$ | | 0.13 | | | | |
|  |  | Variance | Std. Deviation |  |  |  |
| **Model 3** | Random Effect |  |  |  |  |  |
|  | Program | 0.05 | 0.23 |  |  |  |
|  | Research Experience | 0.05 | 0.22 |  |  |  |
|  | Fixed Effect |  |  |  |  |  |
|  | Intercept | 4.72 | 0.13 | 7.24 | 37.49 | <0.0001 |
|  | URE | 0.24 | 0.08 | 420.14 | 2.81 | 0.005 |
| AIC |  | 1203.16 |  |  |  |  |
| $R^2$ | | 0.13 | | | | |

### Values Alignment

We found that students began their UREs with a very high level of values alignment (Model 3: β_0_=5.33, *SE=*0.07, *p*<0.001) and did not, as a group, change in their values alignment from pre to post URE (Model 3: β_1_*=*0.01, *SE=*0.07, *p*=0.856) (see Supplemental Materials). Program did not account for any variance in values alignment, and prior research experience only accounted for 1%, which indicates that the program and prior research experience did not affect students’ values alignment. We observed that students with more than three semesters or summers of prior research experience displayed small but significant gains in values alignment (Model 2: β=0.35, *SE*=0.11, *p*=0.002).

### Perceived Benefits

Similar to the values alignment, we found that students began their UREs at a very high level of perceived benefits (Model 3: β_0_=5.32, *SE=*0.07, *p*<0.001) and did not change in their perceptions of the benefits of doing research from pre to post URE (Model 3: β_1_=-0.06, *SE=*0.06, *p*=0.287) (see Supplemental Materials). Program did not account for any variance in perceived benefits and prior research experience only accounted for 1%. We observed that students with more than three semesters and summers of prior research experience displayed small gains in perceived benefits (Model 2: β=0.025, *SE*=0.10, *p*=0.011). However, the *p* value is greater than our adjusted alpha level of *p*=0.007, indicating a non-significant effect.

### Perceived Costs

We found that students began their UREs reporting a moderate level of perceived costs (Model 3: β_0_=3.53, *SE=*0.14, *p*<0.001). Their perceptions of costs did not change significantly pre- to post-URE (Model 3: β_1_=-0.05, *SE=*0.13, *p*=0.68) (Table 5). In contrast to the other outcomes, we found that program accounted for 23% of variance in students’ perceptions of the cost of research, indicating that the programs in this study differentially affected students’ perceptions of the costs of research. Students’ prior research experience did not account for any variance in their perceptions of the costs of research.

**Table 5.**
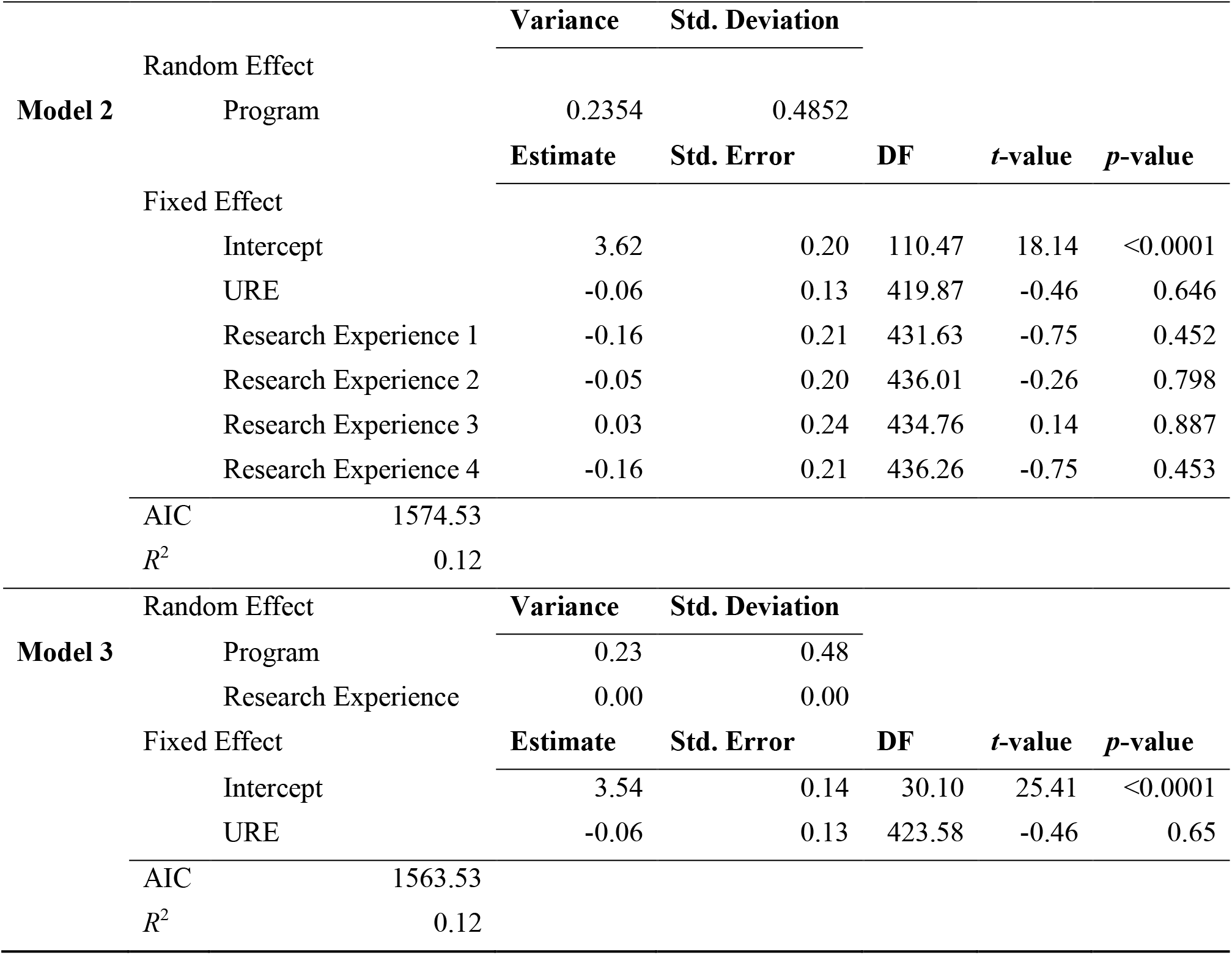
Student perceptions of the cost of doing research vary by program, but not by current or prior research experience. Students reported no change in perceptions of the cost of doing research from pre- to post-URE and no differences in cost perceptions based on their prior research experience. Program had a significant and moderate effect on students’ perceptions of the cost of doing research.

### Graduate School and Career Intentions

On average, we found that students started their remote UREs already intending to attend graduate school (Model 3: β_0_=4.38, *SE=*0.08, *p*<0.001) (see Supplemental Materials). This intention did not change pre- to post-URE (Model 3: β_1_=0.03, *SE=*0.07, *p*=0.717). Likewise, students’ intentions to pursue a career in science were high before completing the program (Model 3: β_0_=4.23, *SE=*0.07, *p*<0.001) and did not change significantly pre- to post-URE (Model 3: β_1_=0.10, *SE=*0.08, *p*=0.196). We found that program accounted for only 2% of variance in graduate school intentions and 1% of variance in career intentions, which suggests that programs did not have different effects on students’ graduate and career intentions.

Similarly, prior research experience accounted for only 1% of variance in graduate school and career intentions, which suggests that different amounts of research experience did not differentially affect students’ intentions. We observed that students with more than three semesters or summers of research experience experienced gains in graduate school intentions (Model 2: β=0.26, SE=0.12, *p*=0.031) and career intentions (β=0.29, *SE*=0.12, *p*=0.019). However, this effect was nonsignificant with our corrected *p*<0.007.

## DISCUSSION

In this study, we first sought to determine whether undergraduates who engage in remote research programs experienced research-related social influence in terms of gains in their self-efficacy, scientific identity, and values alignment (Research Question 1). We found that students in remote UREs experienced outcomes that indicated their integration into the scientific community despite the remote circumstances (Adedokun et al., 2013; Estrada et al., 2011; Robnett et al., 2015). Specifically, students who completed remote UREs experienced significant gains in their scientific self-efficacy, and these gains were largely due to their research experience and not to their particular URE program. Students in remote UREs also experienced gains in their scientific identity, although these gains were more modest than their self-efficacy gains and were related to their specific program. This finding suggests that remote UREs can be productive environments for students’ scientific identity development, but that programs are either attracting or selecting students who differ in their scientific identity or that certain programs are having greater influence on students’ identity development. Students in our study did not experience any changes in the extent to which they perceived their personal values as aligned with the values of the scientific community. Rather, students in this study already felt that their personal values were well-aligned with the values of the scientific community.

We also sought to determine the extent to which students in remote UREs experienced further integration into the scientific community indicated by shifts in their intentions to pursue graduate education and science research-related careers. For the most part, students in this study already intended to pursue a graduate degree and a career in science research, and their intentions did not change significantly from pre- to post-URE. It is encouraging that the challenges of engaging in research remotely did not dissuade students from pursuing graduate school and research careers. Yet, it is also important to note that remote research and perhaps UREs in general may not be a lever for changing students’ plans to pursue graduate education or science research careers because students who seek out or are selected into these programs may already be firm in their intentions.

Although students in this study gained in their scientific self-efficacy, it is worth noting that students started their UREs reporting less confidence in their computational skills. It is unclear whether students’ initial uncertainty about their computational skills is specific to remote research or unique to the last-minute shift from away from bench or field research. As a reminder, most of the students in this study were accepted into their programs *before* decisions were made to offer programs remotely. Regardless, it appears that the programs in this study were able to support students’ development of computational skills.

In keeping with the Expectancy-Value Theory of motivation, we also sought to explore the extent to which undergraduates in remote research programs shifted their perceptions of the benefits and costs of doing research (Research Question 2). Students in this study already perceived high benefits and low costs of research when they started their remote research and their perceptions did not change. Again, it is encouraging that the challenges of engaging in research remotely did not dissuade students from the benefits of research or increase their perceptions of the costs. Interestingly, students’ programs appeared to shape their perceptions of costs of doing research. It may be that some program contexts lessened students’ perceptions of costs and others exacerbated students’ perceptions of costs. The types of institutions that hosted the remote URE programs in our sample varied widely, from masters-granting institutions to high research-intensity universities to non-degree granting research institutes. It may be that differences in research mentor workloads or lifestyles or institutional cultures in these different environments affected students’ perceptions of the costs of doing research. Alternatively, it may be that contextual differences between students’ own undergraduate institutions and the institution that hosted their remote URE program are influencing students’ cost perceptions. Indeed, Duckworth and Yeager (2015) have argued about the importance of considering context dependency of some measures. For instance, a student who has done research at a more teaching-intensive institution and then participates in a summer URE at a highly research-intensive university, or vice versa, may shift substantially in their perceptions of what doing research entails and thus what opportunity costs they might experience if they choose to continue in research.

### How much research experience is enough?

Our results indicate that students with more prior research experience benefited more from remote UREs compared to students with less research experience. Students with the most prior experience reported the most substantial gains in scientific self-efficacy, scientific identity, values alignment, and graduate and career intentions. This finding suggests that students may need at least two or three terms of research experience before they start to realize positive gains from a summer remote URE. There are several plausible explanations for why more experienced researchers realize greater gains in scientific self-efficacy and scientific identity. One possibility is that self-efficacy functions as a positive feedback loop or virtuous cycle. As students gain more experience, they become better at research and are willing to try more things and put forth effort. As a result, they experience more research success and thus become more confident in their abilities to do research. Alternatively, it may be that students who seek out additional research experiences are primed to gain the most. It also may be that students with less research experience do not feel efficacious and thus are less likely to seek out additional research experience, thereby exerting a selection effect. This result provides at least some evidence that, if remote URE programming continues, less experienced students should be prioritized for in-person UREs and more experienced researchers should be prioritized for remote UREs. Alternatively, remote UREs could develop and evaluate additional program elements aimed at better supporting of novice researchers.

### Comparison to In-Person UREs

Overall, we found that students in this study realized scientific self-efficacy growth that resembled the growth observed by Robnett and colleagues (2015) in their longitudinal study of students who completed in-person UREs at colleges and universities across the country. Interestingly, the positive effects observed by Robnett et al. (2015) took place over a period of four semesters of in-person research, while positive effects we observed occurred in a much shorter period – an average of about nine weeks – in entirely remote research. This result may be due to the intensity of the summer experience (∼35-40 hours per week) versus the less intense, more protracted nature of academic year UREs. Alternatively, the remote nature of the programs in this study may have prompted mentors and program leadership to engage more regularly or intentionally with students to ensure they can engage and make progress at a distance. In addition, remote programming may have selected, intentionally or unintentionally, for mentors who were most invested in undergraduate research and undergraduate researchers who were particularly primed to invest time and effort, thereby maximizing the likelihood of students’ experiencing favorable outcomes.

Our results differed to some extent from other longitudinal studies of in-person UREs. Estrada et al. (2018) studied the effects of UREs on the self-efficacy, scientific identity, and values alignment of a cohort of underrepresented minority students in their junior and senior years. Similar to our results, they found that in-person UREs had a small but significant, positive effect on scientific self-efficacy and scientific identity of these more advanced students. In contrast, they found that in-person UREs also had a small but significant, positive effect on students’ values alignment. Hernandez and colleagues (2020) tracked a cohort of STEM students from historically well-represented backgrounds at a research-intensive, public university throughout their four years of college. They also found that in-person research experiences positively predicted scientific self-efficacy and scientific identity but failed to predict values alignment among advanced students, similar to our results. In contrast to our results, Hernandez and colleagues (2020) observed self-efficacy and identity growth among first- and second-year students. It is possible that semester-long (or longer) research experiences are needed to promote these outcomes for less experienced researchers. This would suggest that more experienced researchers are better suited for summer research experiences. Alternatively, the benefits of engaging in undergraduate research early on might not be evident until later in college. As Hernandez and colleagues (2020) note, early social integration through mentorship and research experience exerts a reciprocal longitudinal influence on future engagement with the scientific community.

## LIMITATIONS

There are several limitations of this study that should be considered in interpreting the results. The main limitation is that we designed the study as a single-arm, comparison study; no comparison group of students completing UREs in-person was included because of the circumstances caused by COVID-19. It may be that students who opted to participate in a remote URE were particularly primed for success or that mentors and URE program directors put forth additional effort to ensure as positive experience. It also may be that students were grateful to have any meaningful experience in the midst of the pandemic lockdown and thus responded more favorably than would otherwise be the case. Future research should directly compare remote vs. in-person UREs, ideally using random assignment to one or the other format with students who are willing to do either. Our results provide at least some evidence of the benefits of remote research, which mitigates the ethical concerns associated with such a study.

Another limitation relates to our measure of scientific identity, which demonstrated high internal reliability based on coefficient alpha but suboptimal model fit. Moving forward, researchers should seek to improve this measure by modifying item content and collecting additional validity evidence, including its utility for discriminating among undergraduate students with more or less research experience. More robust frameworks may be needed to better operationalize scientific identity, such as the Carlone and Johnson framework, which conceptualizes scientific identity as a combination of social performance, self-recognition as a “science person,” and knowledge and understanding of science content (Carlone & Johnson, 2006).

Finally, there were limitations related to our sample, which was entirely comprised of biology students. Therefore, our results may be unique to the discipline. Biology research may be more or less amenable to remote research compared to other STEM disciplines. Moreover, as the full extent of the COVID-19 pandemic unfolded, students and mentors who chose to move forward with remote research may possess different personality traits or differing levels of our variables of interest (i.e., scientific identity, scientific self-efficacy) from those who opted out of remote research. Research topics themselves likely changed during the transition to accommodate the remote research arrangement, so researchers who chose to move forward with remote research may have conducted a different type of research than they originally planned on. Lastly, data were collected during a time of social unrest in the United States during summer of 2020. Awareness of social unrest and systematic racism may have affected the well-being of participants, which may have influenced their experience in the remote URE program.

## CONCLUSION

Perhaps the greatest advantage of remote research programs is that they open doors for students who may not have the opportunity to participate in an in-person research program (Erikson *et al*., in press). Remote UREs can allow for more flexible scheduling and enable research participation without the additional costs and logistics of travel and lodging. Thus, remote programs may be a viable method of expanding access to UREs, especially among students who may find it difficult to travel. Although remote UREs have many advantages, their appropriateness should be evaluated on a case-by-case basis and should be considered alongside the advantages and disadvantages of in-person UREs. For example, certain types of research (e.g., computational biology) may be more amenable to remote work. Particular research mentors and undergraduates may be better able to navigate the unstructured nature of remote work.

Certain remote research environments may be more or less accessible for different individuals, such as those who can sit and work on a computer for extended periods of time (Reinholz & Ridgway, 2020). Certain personal situations may make remote research more difficult, such as whether individuals have access to robust internet connections and quiet workspaces (Erikson *et al*., in press). Finally, because students are not able to complete bench work at home, remote UREs may aid in the development of a different skillset than in-person UREs. Thus, students may benefit from completing both types of UREs throughout their undergraduate degree in order to develop a wider variety of skills.

In summary, our work suggests that remote UREs can have a positive effect on student outcomes, but they do not benefit all students equally. The benefits of remote UREs are larger for more experienced researchers compared to less experienced researchers. Given that more experienced researchers benefitted more from remote UREs compared to less experienced researchers, institutions may wish to prioritize selection of less experienced researchers into in-person programs and more experienced researchers into remote or hybrid programs. This would provide less experienced researchers with the supervision and guidance needed to grow while allowing more freedom and flexibility to experienced researchers. Institutions should also consider further developing programming to better meet the needs of novice researchers.

It is important to note that students in this study were *all* conducting their *entire* research experience remotely. In the future, URE programs may wish to consider hybrid designs in which some students are in-person and others are remote, or in which all students participate partly in-person and partly remotely. Students may experience a hybrid program quite differently than a remote program, which could influence their outcomes. We are not aware of any existing research to support the efficacy of a hybrid URE program. If such a program exists, we encourage researchers to investigate differential outcomes for in-person and remote students who are within the same URE program.

## ACKNOWLEDGEMENTS

We thank all of the students, faculty, and other research mentors for their willingness to proceed with remote REU programming and for sharing their experiences so that others could learn. We also thank the Social Psychology of Research Experiences and Education group members for feedback on drafts of this manuscript. This material is based upon work supported by National Science Foundation under Grant No. DBI-2030530. Any opinions, findings, conclusions, or recommendations expressed in this material are those of the authors and do not necessarily reflect the views of any of the funding organizations. The authors dedicate this work to all of the undergraduates seeking to do research and the individuals who provide these opportunities despite challenging circumstances.

