## Supplemental materials for "Virtually the same? Evaluating the effectiveness of remote undergraduate research experiences"

This supplement contains the following:


| Item | Page |
|---|---|
| **Measurement Models - Items and Factor Loadings** | |
| Table S1. Scientific Self-Efficacy | S2 |
| Table S2. Scientific Identity | S3 |
| Table S3. Values Alignment, Benefits, and Costs | S4 |
| Table S4. Higher-Order Factor Loadings | S5 |
| Table S5. Higher-Order Factor Correlations | S6 |
| **Regression Models** | |
| Table S6. Scientific Self-Efficacy | S7 |
| Table S7. Scientific Identity | S7 |
| Table S8. Values Alignment | S8 |
| Table S9. Benefits | S9 |
| Table S10. Costs | S10 |
| Table S11. Graduate School Intentions | S11 |
| Table S12. Career Intentions | S12 |
| **References** | S13 |

### Measurement Models

We present factor loadings from confirmatory factor analysis at Time 1 (before the remote URE) and Time 2 (after the URE) to gain insight into whether students are interpreting the items similarly at both time points. It is noteworthy that factor loadings improve from pre- to post-URE for scientific identity, values alignment, intrinsic 1, intrinsic 2, personal importance, cost, social utility, job utility, and life utility.

**Table S1. Scientific Self-Efficacy Items and Factor Loadings.** This measure of Scientific Self-Efficacy includes 7 published items from Chemers et al. (2011) and Estrada et al. (2011). Items 2 and 6 were authored based on input from the directors of the URE programs included in this study to capture the forms of scientific self-efficacy students would develop during their remote UREs. Response options were: Not confident (1), A little confident (2), Somewhat confident (3), Confident (4), Very confident (5), Extremely confident (6), and I prefer not to respond.

*Please indicate how confident you are in your ability to..*

| Item | Content | Time 1 Factor Loadings | Time 2 Factor Loadings |
|---|---|---|---|
| 1 | Use technical skills (lab or field equipment, instruments, and/or bench or field techniques). | 0.66 | 0.49 |
| 2 | Use computational skills (software, algorithms, and/or quantitative techniques). | 0.45 | 0.47 |
| 3 | Generate a research question to answer. | 0.87 | 0.89 |
| 4 | Develop a hypothesis to test. | 0.84 | 0.90 |
| 5 | Figure out what data/observations to collect and how to collect them. | 0.81 | 0.81 |
| 6 | Trouble-shoot an investigation or experiment. | 0.78 | 0.68 |
| 7 | Create explanations for the results of the study. | 0.81 | 0.76 |
| 8 | Use scientific literature and/or reports to guide research. | 0.68 | 0.72 |
| 9 | Develop theories (integrate and coordinate results from multiple studies). | 0.78 | 0.75 |

**Table S2. Scientific identity items and factor loading.** This measure of Scientific Identity includes 7 published items from Chemers et al. (2011) and Estrada et al. (2011). Response options were: Strongly disagree (1), Moderately disagree (2), Slightly agree (3), Moderately agree (4), Mostly agree (5), Strongly agree (6), and I prefer not to respond (7). Response options were positively weighted to avoid a ceiling effect.

*Please indicate the extent to which you agree with the following statements.*

| Item | Content | Time 1 Factor Loadings | Time 2 Factor Loadings |
|---|---|---|---|
| 1 | I have a strong sense of belonging to the community of scientists. | 0.52 | 0.67 |
| 2 | I derive great personal satisfaction from working on a team of scientists. | 0.59 | 0.75 |
| 3 | I think of myself as a scientist. | 0.62 | 0.78 |
| 4 | The daily work of a scientist is appealing to me. | 0.64 | 0.78 |
| 5 | I feel like I belong in the field of science. | 0.66 | 0.85 |
| 6 | In general, being a scientist is an important part of my self-image. | 0.86 | 0.86 |
| 7 | Being a scientist is an important reflection of who I am. | 0.88 | 0.88 |

**Table S3. Values alignment, benefit, and cost items and lower-order factor loadings.** We used the Values Alignment measure published in Estrada et al., 2011. Response options were: Not like me (1), A little like me (2), Somewhat like me (3), Like me (4), Very much like me (5), Extremely like me (6), and I prefer not to respond (7). To measure Intrinsic Value, Personal Importance, Social Utility, Job Utility, Life Utility, and Cost, we adapted measures from Gaspard et al. (2015) measures by changing "math" to "research." Response options were: Strongly disagree (1), Moderately disagree (2), Slightly agree (3), Moderately agree (4), Mostly agree (5), Strongly agree (6), and I prefer not to respond (7). Response options were positively weighted to avoid a ceiling effect.

*Please indicate the extent to which you agree with the following statements.*

| Factor | | Item | T1 Factor Loadings | T2 Factor Loadings |
|---|---|---|---|---|
| **Higher-Order Factor 1: Values Alignment** | | | | |
| Values Alignment | 1 | A person who thinks it is valuable to conduct research that builds the world's scientific knowledge. | 0.68 | 0.85 |
| | 2 | A person who feels discovering something new in the sciences is thrilling. | 0.83 | 0.88 |
| | 3 | A person who thinks discussing new theories and ideas between scientists is important. | 0.79 | 0.81 |
| | 4 | A person who thinks that scientific research can solve many of today's world challenges. | 0.59 | 0.69 |
| **Higher-Order Factor 2: Benefits** | | | | |
| Intrinsic Value 1 | 5 | Research is fun to me. | 0.91 | 0.97 |
| | 6 | I like doing research. | 0.97 | 0.99 |
| | 7 | I enjoy dealing with research topics. | 0.87 | 0.92 |
| Intrinsic Value 2 | 8 | It is important to me to be good at research. | 0.79 | 0.94 |
| | 9 | Being good at research means a lot to me. | 0.90 | 0.97 |
| | 10 | Performing well in research is important to me. | 0.76 | 0.89 |
| Personal Importance | 11 | I care a lot about remembering things I learn when conducting research. | 0.64 | 0.65 |
| | 12 | I'm really keen on learning a lot about research. | 0.74 | 0.80 |
| | 13 | Research is very important to me personally. | 0.85 | 0.91 |
| Social Utility | 17 | Being well versed in research will prepare me to help my community. | 0.66 | 0.82 |
| | 18 | I can do good in the world based on my knowledge of research. | 0.74 | 0.82 |
| | 19 | If I know a lot about research, I can make a difference in the world. | 0.67 | 0.83 |
| Job Utility | 20 | Doing well in research will improve my chances of finding a job after college. | 0.76 | 0.80 |
| | 21 | The skills I develop in research will help me be successful in my career. | 0.80 | 0.80 |
| | 22 | Learning how to conduct research is worthwhile because it improves my career prospects. | 0.86 | 0.88 |
| Life Utility | 23 | Research will help me in life. | 0.74 | 0.82 |
| | 24 | I will often need research in my life. | 0.82 | 0.90 |
| | 25 | Research comes in handy in everyday life. | 0.67 | 0.78 |

**Table S3. Values alignment, benefit, and cost items and lower-order factor loadings. (continued)**

*Please indicate the extent to which you agree with the following statements.*

| Factor | | Item | T1 Factor Loadings | T2 Factor Loadings |
|---|---|---|---|---|
| **Higher-Order Factor 3: Cost** | | | | |
| Cost | 14 | I have to give up other activities that I like to be successful at research. | 0.79 | 0.90 |
| | 15 | I have to give up a lot to do well in research. | 0.96 | 0.93 |
| | 16 | I'd have to sacrifice a lot of free time to be good at research. | 0.73 | 0.81 |

**Table S4. Values alignment, benefit, and Cost higher-order factor loadings.** Loadings for higher-order factors with only one lower-order construct (i.e., Alignment, Cost) will always be 1.00 and are not meaningful. Please see Figure 1 in the main manuscript for pre-URE higher-order factor loadings

| | Post-URE |
|---|---|
| **Higher-Order Factor 1: Alignment** | |
| Alignment | 1.00 |
| **Higher-Order Factor 2: Benefits** | |
| Intrinsic 1 | 0.77 |
| Intrinsic 2 | 0.81 |
| Personal Importance | 0.97 |
| Social Utility | 0.78 |
| Job Utility | 0.68 |
| Life Utility | 0.79 |
| **Higher-Order Factor 3: Cost** | |
| Cost | 1.00 |

**Table S5. Higher-order factor correlations.** Factor correlations are reported for Time 2 (post-URE).

| | Alignment | Intrinsic 1 | Intrinsic 2 | Personal Importance | Cost | Social Utility | Job Utility | Life Utility |
|---|---|---|---|---|---|---|---|---|
| Alignment | | | | | | | | |
| Intrinsic 1 | 0.469 | | | | | | | |
| Intrinsic 2 | 0.459 | 0.594 | | | | | | |
| Personal Importance | 0.466 | 0.814 | 0.786 | | | | | |
| Cost | -0.101 | -0.021 | 0.081 | 0.139 | | | | |
| Social Utility | 0.314 | 0.282 | 0.366 | 0.473 | 0.098 | | | |
| Job Utility | 0.210 | 0.172 | 0.468 | 0.393 | 0.169 | 0.451 | | |
| Life Utility | 0.362 | 0.496 | 0.716 | 0.739 | 0.226 | 0.607 | 0.619 | |

### Regression Models

**Table S6. Students do not differ on scientific self-efficacy by program.** In Model 1, Program only accounted for 4% of students' scientific self-efficacy. See Table 3 in the main manuscript for Model 2 and Model 3, which include a fixed effect of students' prior research experience and a random effect of students' prior research experience, respectively.

| | | | | | | | |
|---|---|---|---|---|---|---|---|
| | | | **Variance** | **Std. Deviation** | | | |
| **Model 1** | Random Effect | | | | | | |
| | Program | | 0.04 | 0.20 | | | |
| | | | **Estimate** | **Std. Error** | **DF** | ***t*-value** | ***p*-value** |
| | Fixed Effect | | | | | | |
| | Intercept | | 3.62 | 0.07 | 35.12 | 49.07 | <.0001 |
| | URE | | 0.64 | 0.08 | 422.20 | 7.82 | <.0001 |
| | AIC | 1165.448 | | | | | |
| | $R^2$ | 0.1604985 | | | | | |

**Table S7. Students do not differ on scientific identity by program.** In Model 1, Program only accounted for 4% of variance in students' scientific identity. See Table 4 in the main manuscript for Model 2 and Model 3 which include a fixed effect of students' prior research experience and a random effect of students' prior research experience, respectively.

| | | | | | | | |
|---|---|---|---|---|---|---|---|
| | | | **Variance** | **Std. Deviation** | | | |
| **Model 1** | Random Effect | | | | | | |
| | Program | | 0.04 | 0.20 | | | |
| | | | **Estimate** | **Std. Error** | **DF** | ***t*-value** | ***p*-value** |
| | Fixed Effect | | | | | | |
| | Intercept | | 4.71 | 0.08 | 42.19 | 61.56 | <.0001 |
| | URE | | 0.24 | 0.09 | 427.99 | 2.75 | 0.01 |
| | AIC | 1217.385 | | | | | |
| | $R^2$ | 0.06 | | | | | |

**Table S8. More experienced researchers see stronger gains in values alignment compared to less experienced researchers.** For most students, there was no effect of the remote research experience or their program on values alignment. There was a small yet significant effect for students with more than three semesters of prior research experience.

| | | | | | | | |
|---|---|---|---|---|---|---|---|
| **Model 1** | | | **Variance** | **Std. Deviation** | | | |
| | Random Effect | | | | | | |
| | Program | | 0.00 | 0.05 | | | |
| | | | **Estimate** | **Std. Error** | **DF** | ***t*-value** | ***p*-value** |
| | Fixed Effect | | | | | | |
| | Intercept | | 5.32 | 0.05 | 50.24 | 106.73 | <.0001 |
| | URE | | 0.02 | 0.07 | 431.87 | 0.25 | 0.800 |
| | AIC | 1010.45 | | | | | |
| | $R^2$ | 0.01 | | | | | |
| **Model 2** | | | **Variance** | **Std. Deviation** | | | |
| | Random Effect | | | | | | |
| | Program | | 0.01 | 0.08 | | | |
| | | | **Estimate** | **Std. Error** | **DF** | ***t*-value** | ***p*-value** |
| | Fixed Effect | | | | | | |
| | Intercept | | 5.22 | 0.09 | 227.09 | 57.12 | <.0001 |
| | URE | | 0.01 | 0.07 | 422.95 | 0.18 | 0.858 |
| | Research Experience 1 | | 0.02 | 0.11 | 443.86 | 0.16 | 0.870 |
| | Research Experience 2 | | 0.08 | 0.11 | 443.39 | 0.78 | 0.434 |
| | Research Experience 3 | | 0.08 | 0.12 | 441.85 | 0.62 | 0.534 |
| | Research Experience 4 | | 0.35 | 0.11 | 442.19 | 3.16 | 0.002 |
| | AIC | 1012.53 | | | | | |
| | $R^2$ | 0.04 | | | | | |
| **Model 3** | Random Effect | | **Variance** | **Std. Deviation** | | | |
| | Program | | 0.00 | 0.07 | | | |
| | Research Experience | | 0.01 | 0.12 | | | |
| | Fixed Effect | | **Estimate** | **Std. Error** | **DF** | ***t*-value** | ***p*-value** |
| | Intercept | | 5.33 | 0.07 | 6.82 | 71.50 | <.0001 |
| | URE | | 0.01 | 0.07 | 422.43 | 0.18 | 0.856 |
| | AIC | 1003.95 | | | | | |
| | $R^2$ | 0.04 | | | | | |

**Table S9. Students' perceptions of the benefits of doing research do not change over the course of the URE.** Students enter their remote URE programs with highly positive perceptions of the benefits of research, and those perceptions do not change over the course of the remote summer URE.

| | | | | | | | |
|---|---|---|---|---|---|---|---|
| | | | **Variance** | **Std. Deviation** | | | |
| | Random Effect | | | | | | |
| | Program | | 0.00 | 0.07 | | | |
| | | | **Estimate** | **Std. Error** | **DF** | ***t*-value** | ***p*-value** |
| **Model 1** | Fixed Effect | | | | | | |
| | Intercept | | 5.32 | 0.05 | 30.65 | 116.46 | <.0001 |
| | URE | | -0.06 | 0.06 | 415.53 | -1.06 | 0.291 |
| | AIC | 887.83 | | | | | |
| | $R^2$ | 0.01 | | | | | |
| | | | **Variance** | **Std. Deviation** | | | |
| | Random Effect | | | | | | |
| | Program | | 0.01 | 0.10 | | | |
| | | | **Estimate** | **Std. Error** | **DF** | ***t*-value** | ***p*-value** |
| | Fixed Effect | | | | | | |
| | Intercept | | 5.26 | 0.08 | 192.20 | 63.66 | <.0001 |
| **Model 2** | URE | | -0.06 | 0.06 | 414.00 | -1.06 | 0.288 |
| | Research Experience 1 | | -0.09 | 0.10 | 438.00 | -0.98 | 0.328 |
| | Research Experience 2 | | 0.00 | 0.09 | 440.00 | 0.00 | 1.000 |
| | Research Experience 3 | | 0.12 | 0.11 | 440.00 | 1.10 | 0.272 |
| | Research Experience 4 | | 0.25 | 0.10 | 440.00 | 2.55 | 0.011 |
| | AIC | 888.92 | | | | | |
| | $R^2$ | 0.06 | | | | | |
| | Random Effect | | **Variance** | **Std. Deviation** | | | |
| | Program | | 0.00 | 0.09 | | | |
| | Research Experience | | 0.01 | 0.12 | | | |
| **Model 3** | Fixed Effect | | **Estimate** | **Std. Error** | **DF** | ***t*-value** | ***p*-value** |
| | Intercept | | 5.32 | 0.07 | 7.04 | 75.24 | <.0001 |
| | URE | | -0.06 | 0.06 | 411.72 | -1.07 | 0.287 |
| | AIC | 879.8508 | | | | | |
| | $R^2$ | 0.04 | | | | | |

**Table S10. Students' perceptions of the cost of doing research vary by program.** In Model 1, Program accounted for 22% of variance in students' perceptions of the cost of doing research. See Table 5 in the Manuscript for Model 2 and Model 3, which include a fixed effect of students' prior research experience and a random effect of students' prior research experience, respectively.

| | | | | | | | |
|---|---|---|---|---|---|---|---|
| | | | **Variance** | **Std. Deviation** | | | |
| | Random Effect | | | | | | |
| **Model 1** | Program | | 0.22 | 0.47 | | | |
| | | | **Estimate** | **Std. Error** | **DF** | ***t*-value** | ***p*-value** |
| | Fixed Effect | | | | | | |
| | Intercept | | 3.53 | 0.14 | 30.93 | 25.80 | <.0001 |
| | URE | | -0.05 | 0.13 | 425.86 | -0.41 | 0.680 |
| | AIC | 1568.33 | | | | | |
| | $R^2$ | 0.11 | | | | | |

**Table S11. Students' intentions to pursue graduate education do not change over the course of the URE.** Most students enter their remote URE programs intending to pursue graduate school, and that intention does not change over the course of the remote summer URE.

| | | | | | | | |
|---|---|---|---|---|---|---|---|
| **Model 1** | | | **Variance** | **Std. Deviation** | | | |
| | Random Effect | | | | | | |
| | Program | | 0.02 | 0.14 | | | |
| | | | **Estimate** | **Std. Error** | **DF** | ***t*-value** | ***p*-value** |
| | Fixed Effect | | | | | | |
| | Intercept | | 4.38 | 0.06 | 52.88 | 71.18 | <.0001 |
| | URE | | 0.03 | 0.07 | 436.03 | 0.36 | 0.720 |
| | AIC | 1091.778 | | | | | |
| | $R^2$ | 0.03 | | | | | |
| **Model 2** | | | **Variance** | **Std. Deviation** | | | |
| | Random Effect | | | | | | |
| | Program | | 0.01 | 0.12 | | | |
| | | | **Estimate** | **Std. Error** | **DF** | ***t*-value** | ***p*-value** |
| | Fixed Effect | | | | | | |
| | Intercept | | 4.28 | 0.10 | 252.22 | 42.13 | <.0001 |
| | URE | | 0.03 | 0.07 | 430.59 | 0.36 | 0.718 |
| | Research Experience 1 | | -0.08 | 0.12 | 445.11 | -0.72 | 0.475 |
| | Research Experience 2 | | 0.13 | 0.12 | 446.00 | 1.10 | 0.274 |
| | Research Experience 3 | | 0.20 | 0.14 | 445.92 | 1.48 | 0.139 |
| | Research Experience 4 | | 0.26 | 0.12 | 445.84 | 2.17 | 0.031 |
| | AIC | 1095.039 | | | | | |
| | $R^2$ | 0.04972416 | | | | | |
| **Model 3** | Random Effect | | **Variance** | **Std. Deviation** | | | |
| | Program | | 0.02 | 0.13 | | | |
| | Research Experience | | 0.01 | 0.12 | | | |
| | Fixed Effect | | **Estimate** | **Std. Error** | **DF** | ***t*-value** | ***p*-value** |
| | Intercept | | 4.38 | 0.08 | 9.24 | 54.93 | <.0001 |
| | URE | | 0.03 | 0.07 | 431.67 | 0.36 | 0.717 |
| | AIC | 1086.484 | | | | | |
| | $R^2$ | 0.05 | | | | | |

**Table S12. Students' intentions to pursue a research career do not change over the course of the URE.** Most students enter their remote URE programs intending to pursue a career in research, and that intention does not change over the course of the remote summer URE.

| | | | | | | | |
|---|---|---|---|---|---|---|---|
| **Model 1** | | | **Variance** | **Std. Deviation** | | | |
| | Random Effect | | | | | | |
| | Program | | 0.01 | 0.10 | | | |
| | | | **Estimate** | **Std. Error** | **DF** | ***t*-value** | ***p*-value** |
| | Fixed Effect | | | | | | |
| | Intercept | | 4.22 | 0.06 | 72.18 | 72.76 | <.0001 |
| | URE | | 0.10 | 0.08 | 439.78 | 1.29 | 0.20 |
| | AIC | 1109.168 | | | | | |
| | $R^2$ | 0.02 | | | | | |
| **Model 2** | | | **Variance** | **Std. Deviation** | | | |
| | Random Effect | | | | | | |
| | Program | | 0.01 | 0.10 | | | |
| | | | **Estimate** | **Std. Error** | **DF** | ***t*-value** | ***p*-value** |
| | Fixed Effect | | | | | | |
| | Intercept | | 4.10 | 0.10 | 284.57 | 40.04 | <.0001 |
| | URE | | 0.10 | 0.08 | 433.36 | 1.29 | 0.196 |
| | Research Experience 1 | | 0.06 | 0.12 | 445.82 | 0.50 | 0.619 |
| | Research Experience 2 | | 0.06 | 0.12 | 445.79 | 0.52 | 0.607 |
| | Research Experience 3 | | 0.25 | 0.14 | 445.17 | 1.79 | 0.074 |
| | Research Experience 4 | | 0.29 | 0.12 | 445.23 | 2.36 | 0.019 |
| | AIC | 1114.60 | | | | | |
| | $R^2$ | 0.04 | | | | | |
| **Model 3** | Random Effect | | **Variance** | **Std. Deviation** | | | |
| | Program | | 0.01 | 0.10 | | | |
| | Research Experience | | 0.01 | 0.09 | | | |
| | Fixed Effect | | **Estimate** | **Std. Error** | **DF** | ***t*-value** | ***p*-value** |
| | Intercept | | 4.23 | 0.07 | 9.10 | 59.00 | <.0001 |
| | URE | | 0.10 | 0.08 | 433.54 | 1.29 | 0.196 |
| | AIC | 1105.15 | | | | | |
| | $R^2$ | 0.03 | | | | | |
